## Supplementary Figures for "Relative enrichment – a density-based colocalization measure for single-molecule localization microscopy"

Ejdrup et al.

**Supplementary information**

Supplementary Figures 1-3

**Supplementary Figure 1.**

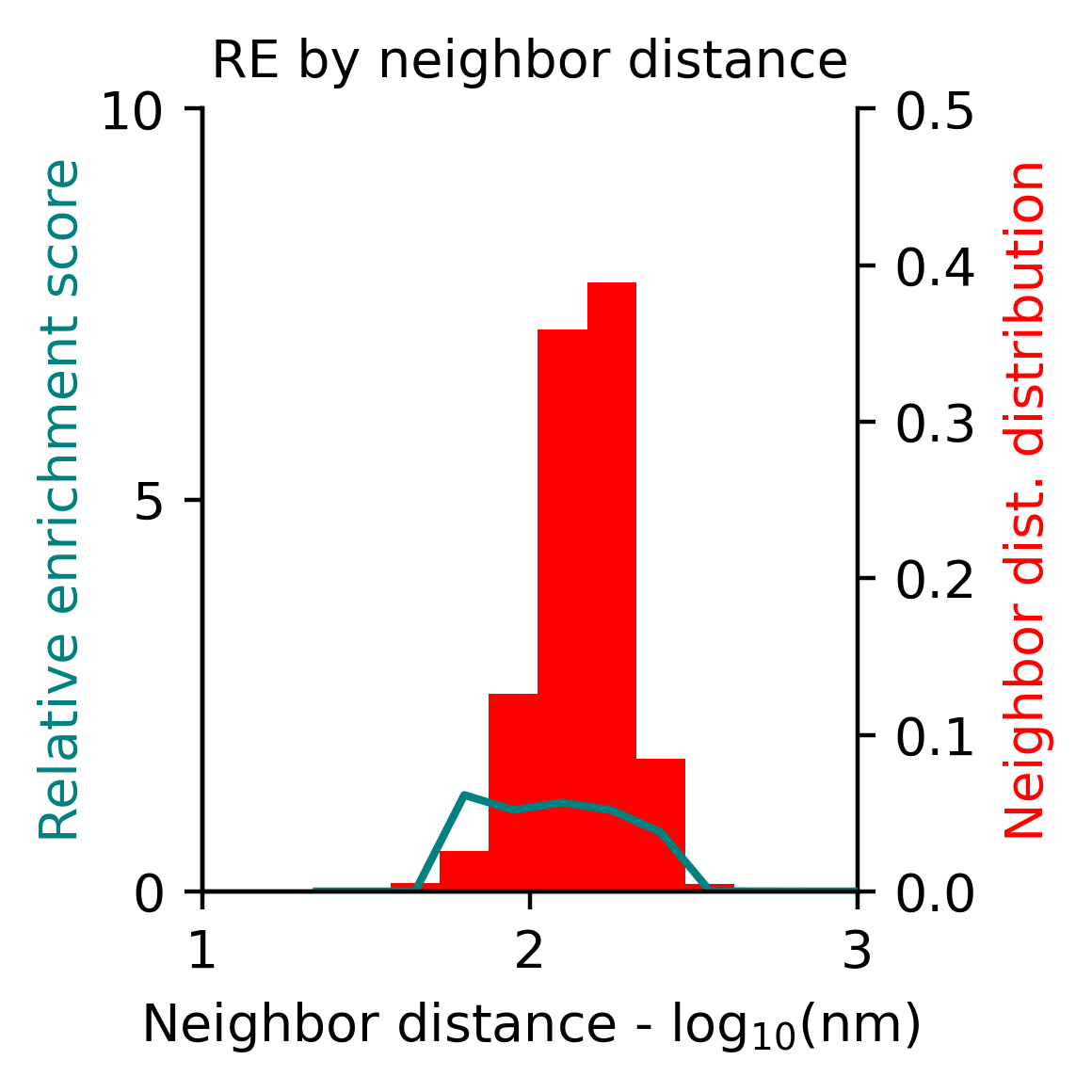

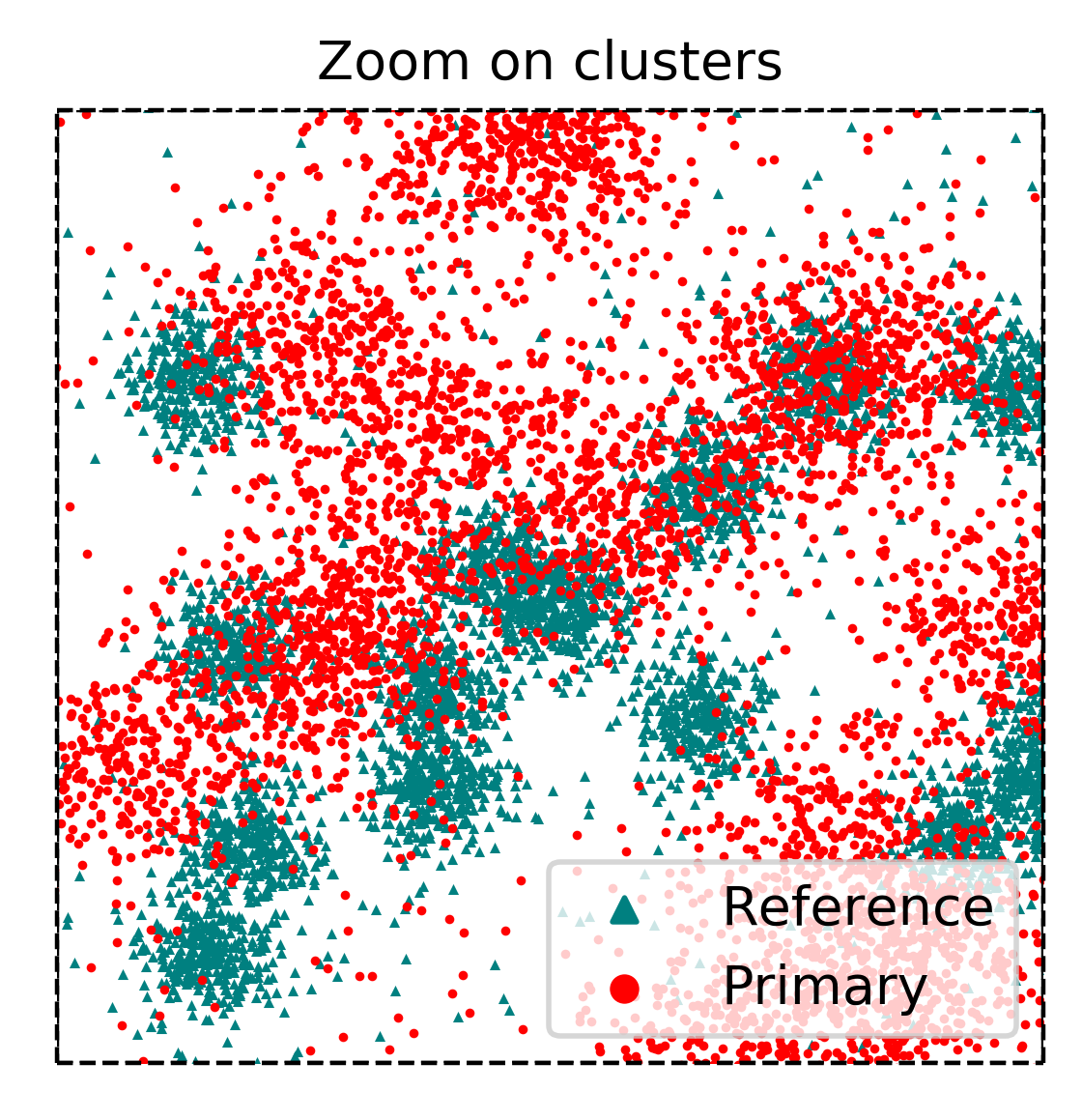

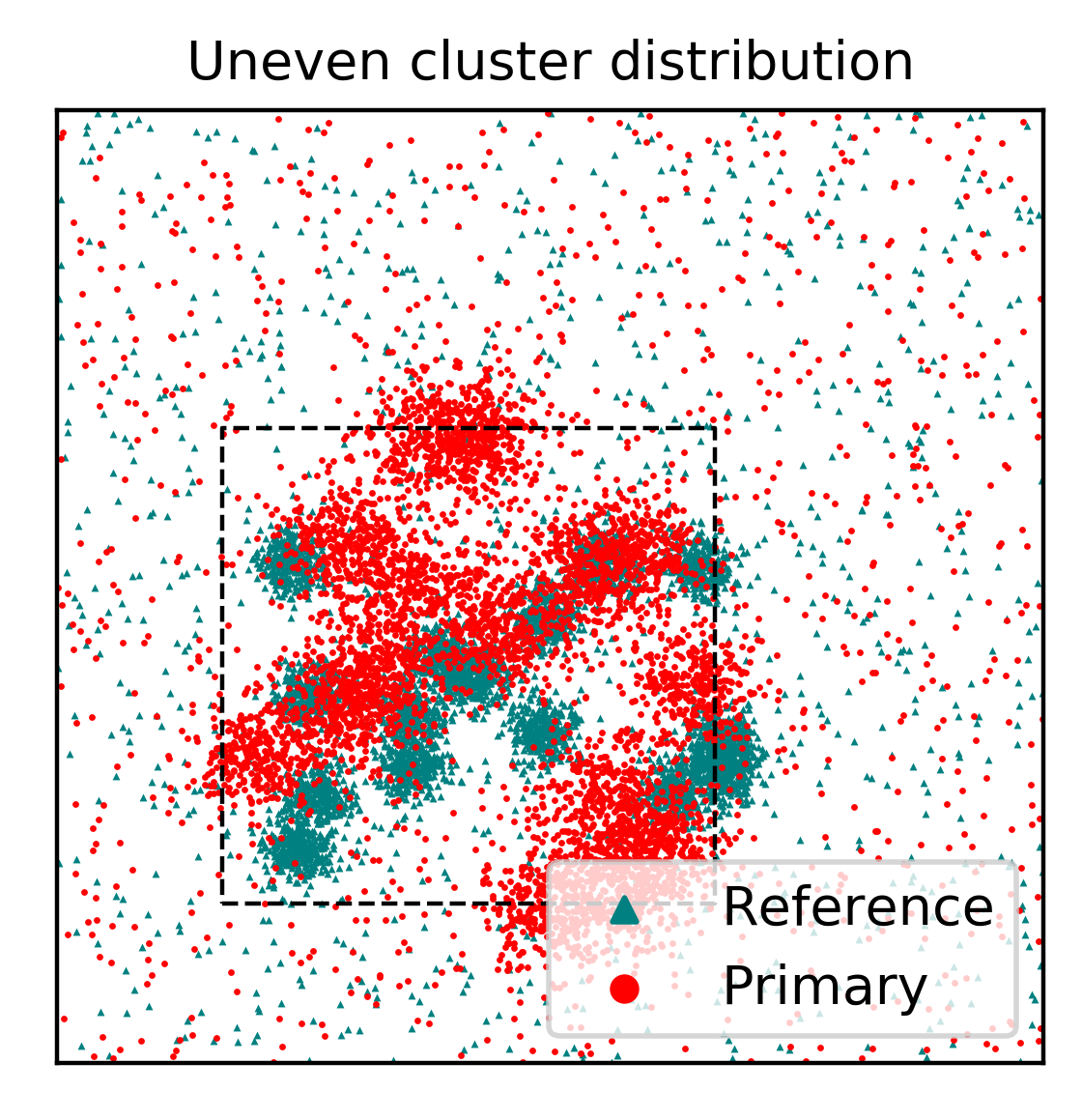

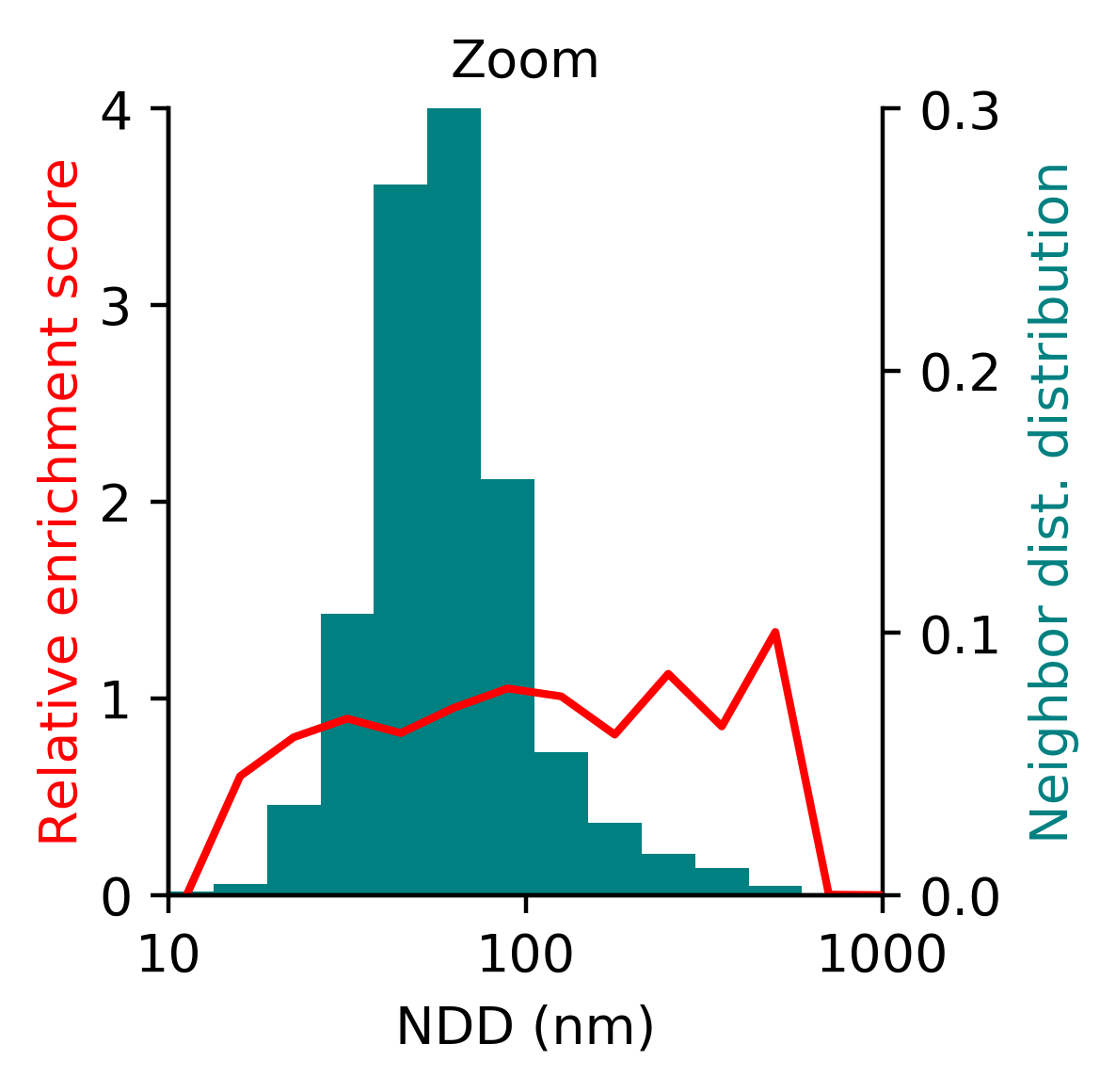

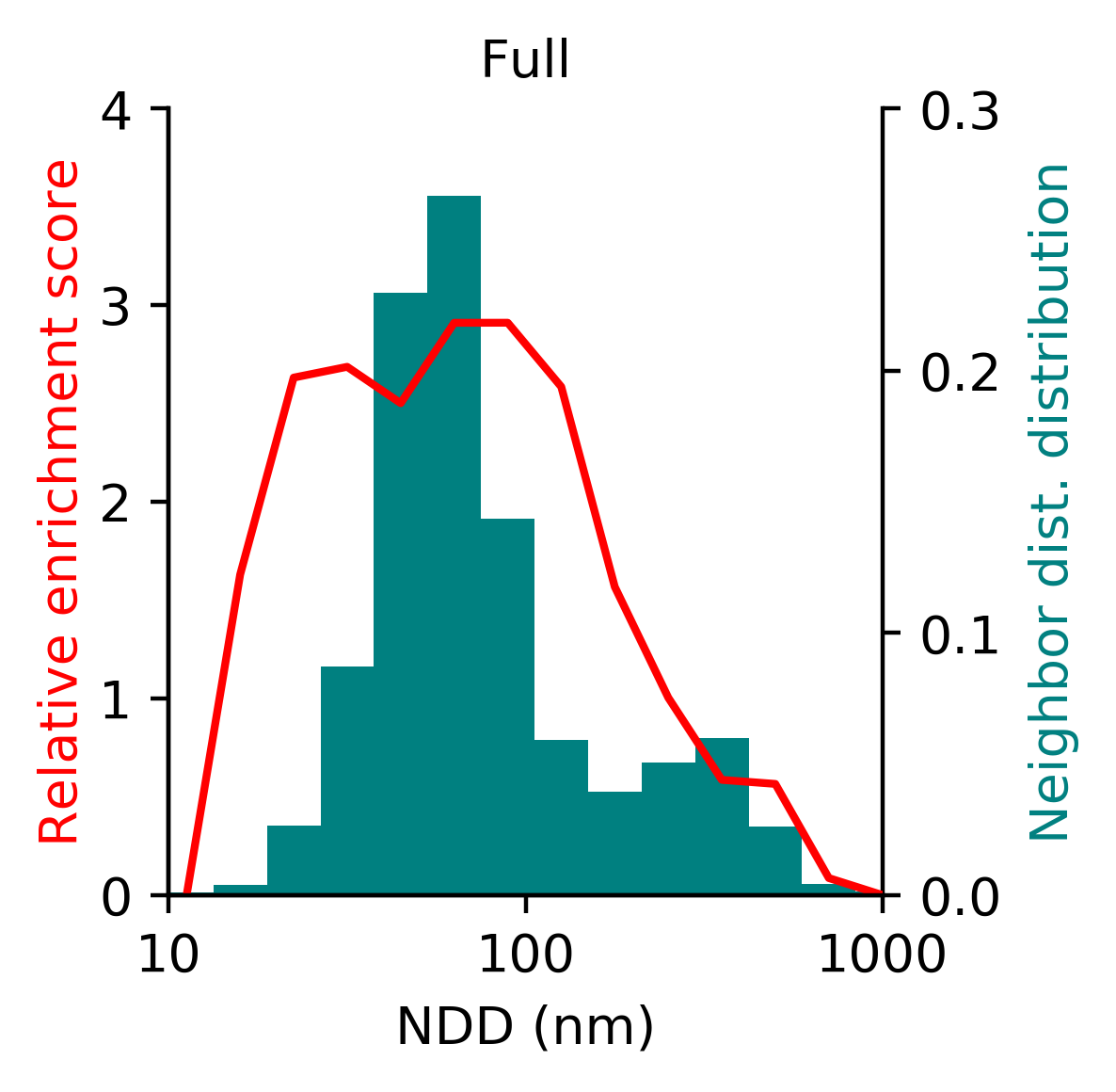

A

B

D

E

C

**Supporting Figure 1. Uneven cluster distribution.**
(A) Mean RE score across reference densities, binned by nearest neighbour distance (NND), for data on Fig. 2G, but with the species reversed. The narrow reference histogram reflects the uniform distribution.
(B) Simulation of two species, with a clustered fraction in both species confined to the same area.
(C) Mean RE score across reference densities of the entire area in (B).
(D) Zoom on the dashed area in (B).
(E) Mean RE score across reference densities of the zoom on (E).

**Supplementary Figure 2.**

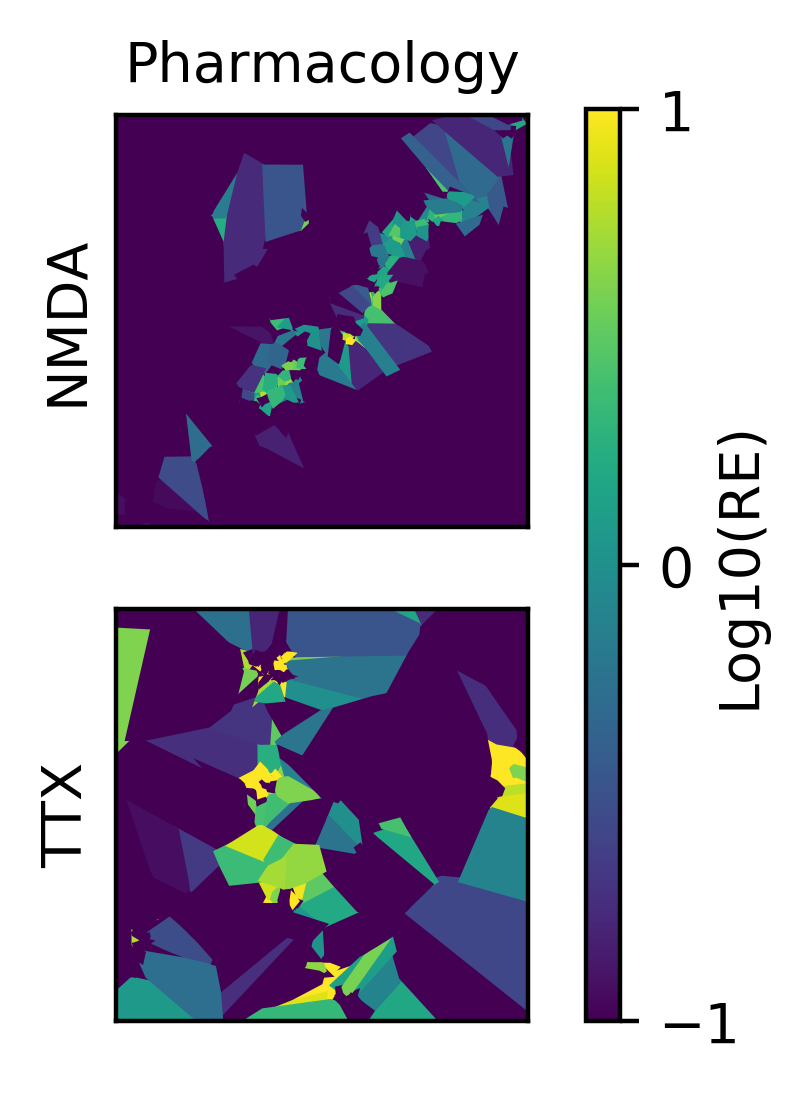

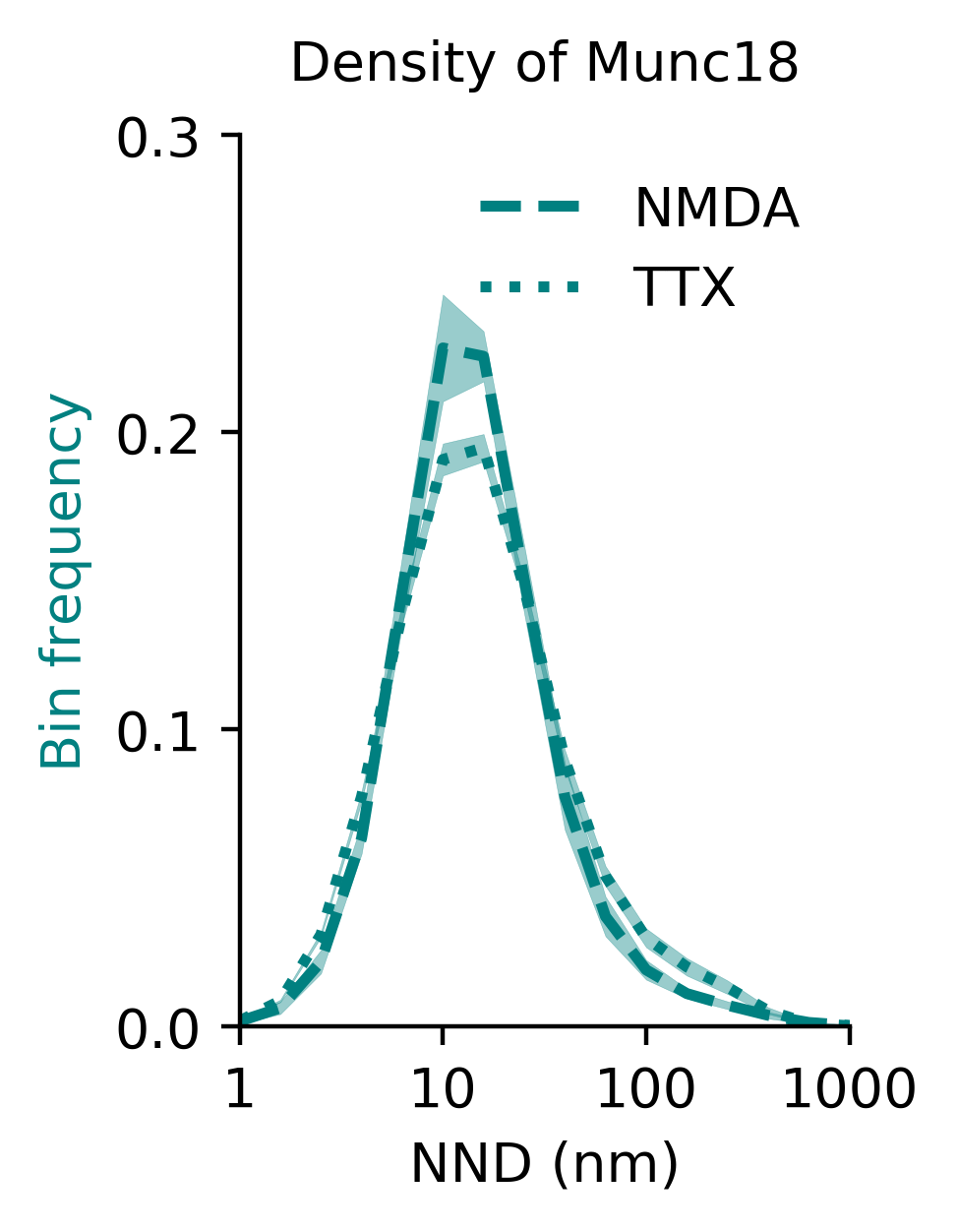

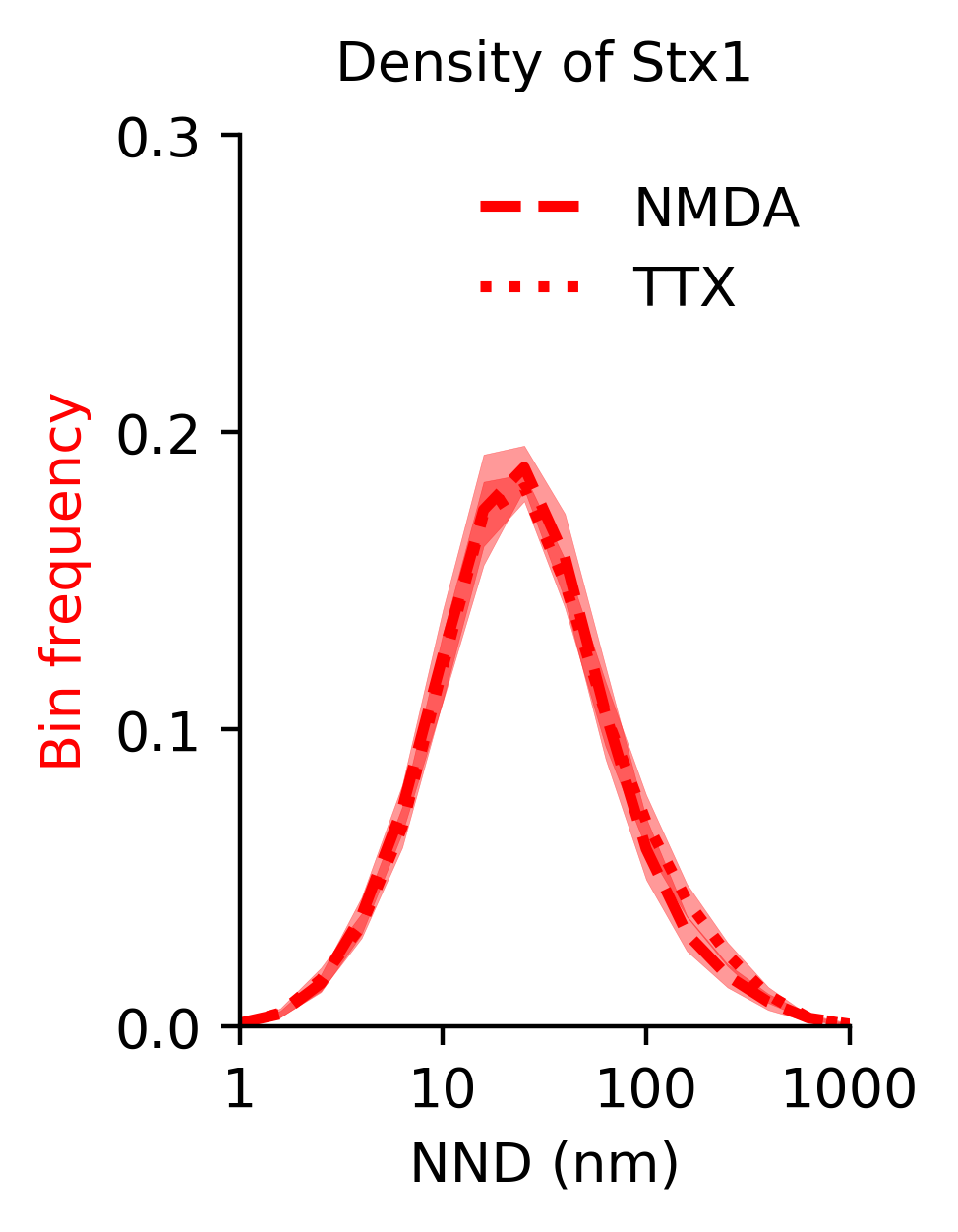

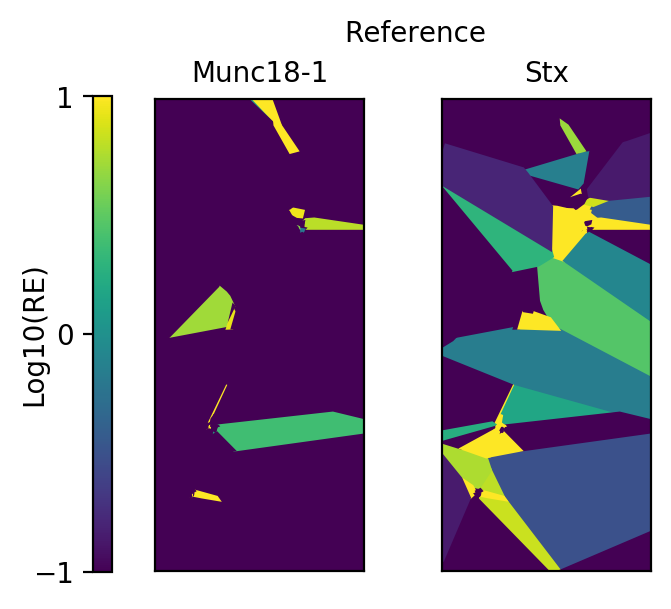

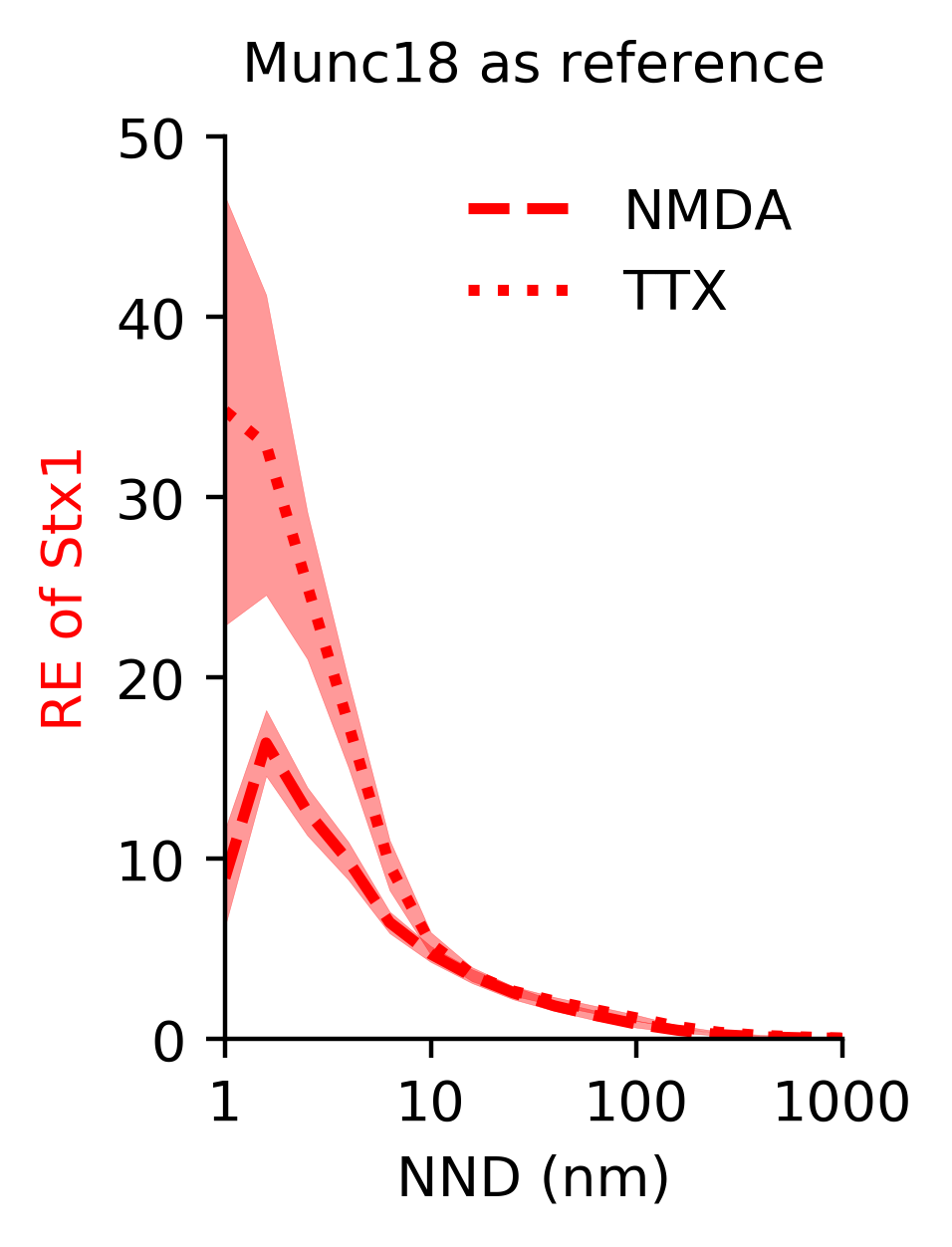

A

D

B

E

C

**Supporting Figure 2. Munc18-1 and Stx1 extra figures.**
(A) Voronoï regions of Fig. 3C color-coded for RE values.
(B) Voronoï regions of Fig. 3G color-coded for RE values.
(C) Density of Stx1 after NMDA (dashed, n = 7 images) or TTX (dotted, n = 8 images) treatment. Shaded area indicates S.E.M..
(D) Relative enrichment of Stx1 across Munc18-1 densities after NMDA (dashed, n = 7 images) or TTX (dotted, n = 8 images) treatment. Shaded area indicates S.E.M..
(E) Density of Munc18-1 after NMDA (dashed, n = 7 images) or TTX (dotted, n = 8 images) treatment. Shaded area indicates S.E.M..

**Supplementary Figure 3.**

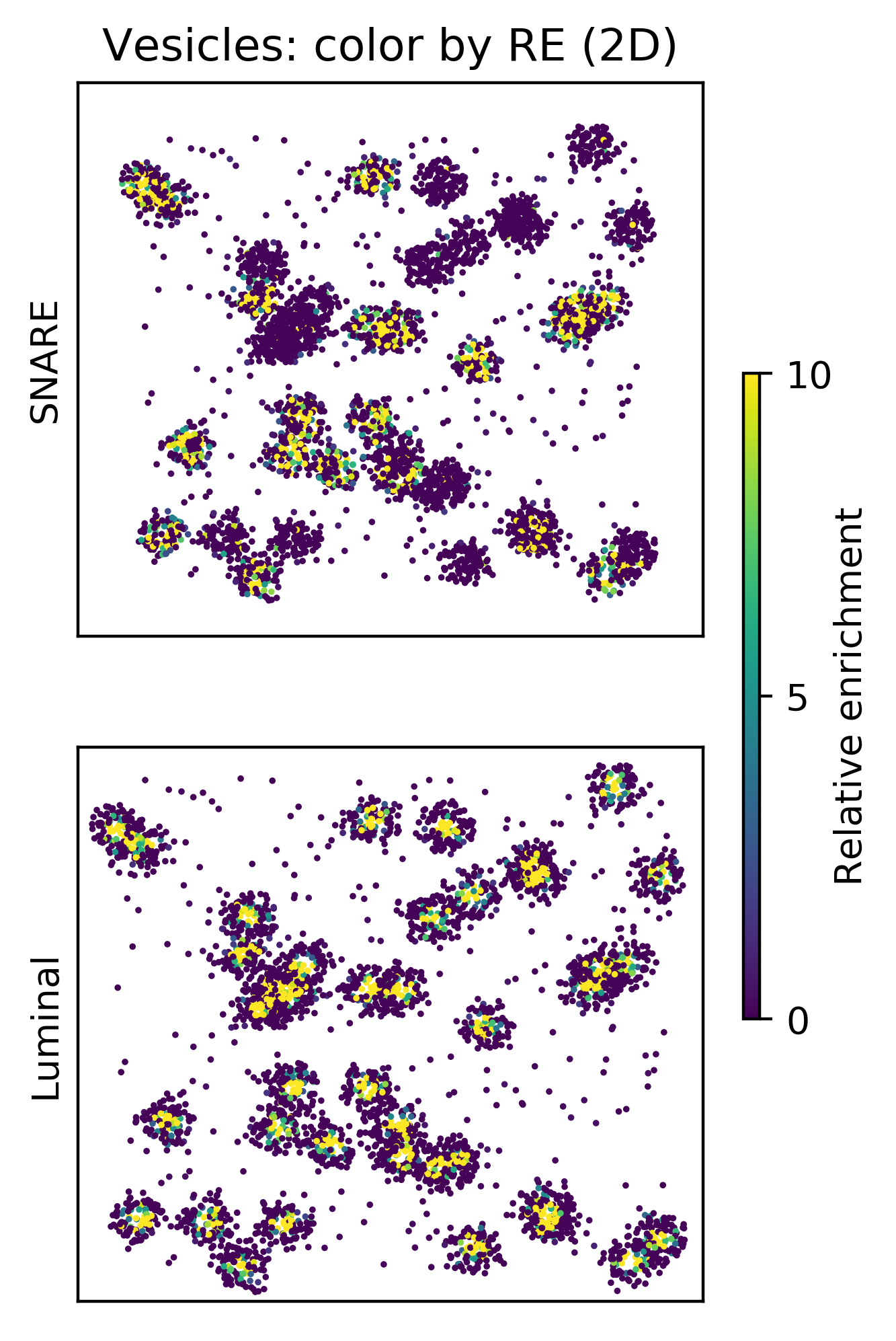

A

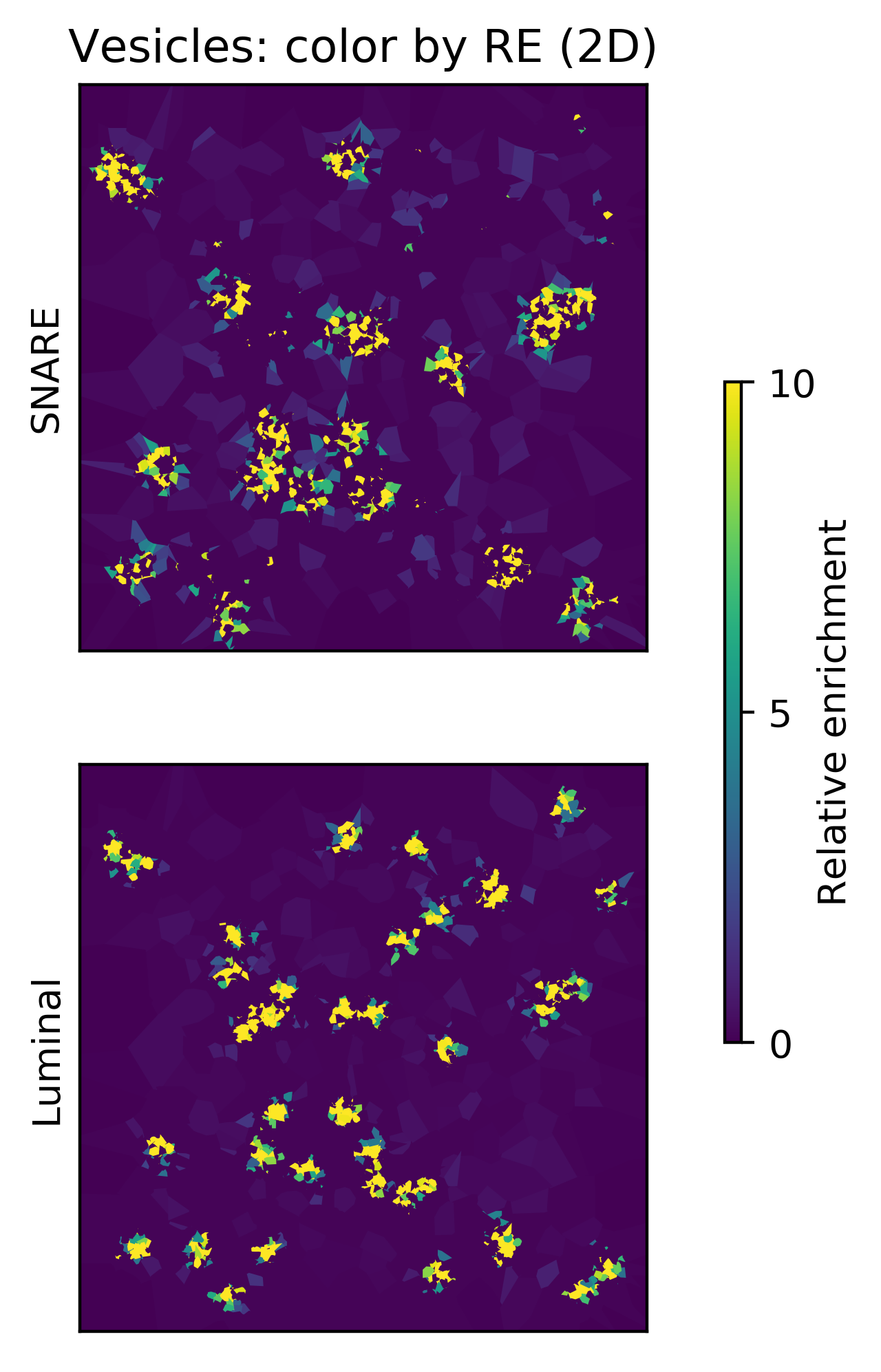

B

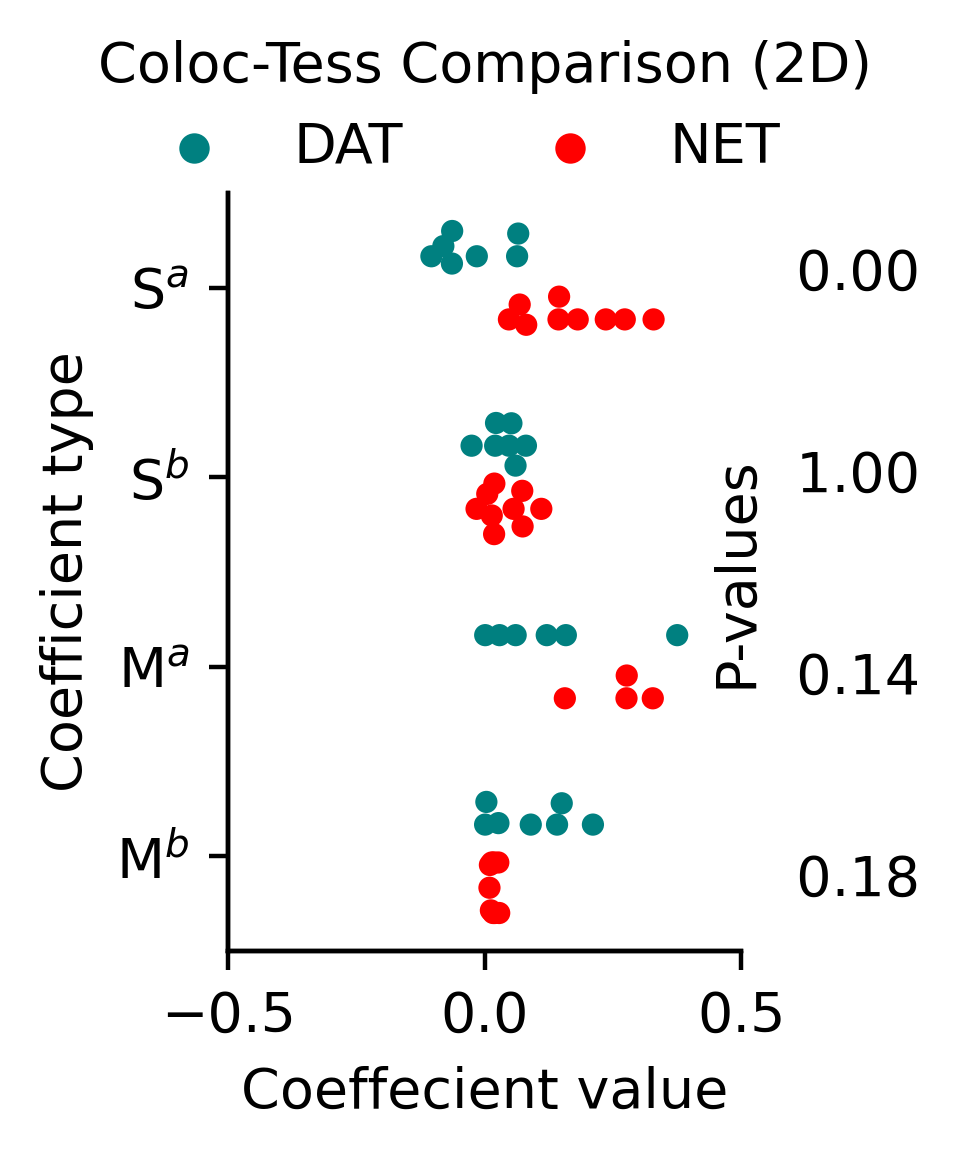

C

**Supporting Figure 3. Color-coding of simulated vesicular species.**
(A) Individual localization of the vesicle species color-coded for RE score of either the AZ (top) or cargo (bottom) species.
(B) Same as (A), but with color-coded Voronoï regions.
(C) Spearman (S) and Mander’s (M) coefficient computed in 2D as per [12] (H_0_: DAT = NET. S^EEA1^, p = 0.00; S^D/N^, p = 1.00; M^EEA1^, p = 0.15; M^D/N^, p = 0.18. Unpaired, one-sided student’s t-test by image, FWER correction with Bonferroni-Holm).
